# Identification of the role of SEL1L in platelet function through a multi-species genetic investigation

**DOI:** 10.1101/2025.04.30.651507

**Authors:** Anna R. Dahlgren, Francesca Careddu, Jeffrey W. Norris, Reheman Adili, Erin M. Kropp, Qing Li, Michael Holinstat, Ida Biunno, Alessandra Balduini, Fern Tablin, Jordan A. Shavit, Carrie J. Finno

**Affiliations:** Department of Population Health and Reproduction, School of Veterinary Medicine, University of California Davis, Davis, CA; Department of Pediatrics, University of Michigan, Ann Arbor, MI; Department of Molecular Medicine, University of Pavia, Pavia, Italy; Department of Anatomy, Physiology and Cell Biology, School of Veterinary Medicine, University of California Davis, Davis, CA; Department of Pharmacology, Midwestern University, Glendale, AZ; Blood Works Northwest Research Institute, Seattle, WA; Department of Internal Medicine, Hematology Oncology Division, University of Michigan-Ann Arbor, Ann Arbor, MI; VA Ann Arbor Healthcare System, Medicine Service (111), Ann Arbor, MI; Department of Cell and Developmental Biology, University of Michigan, Ann Arbor, MI; Department of Pharmacology, University of Michigan, Ann Arbor, MI; Department of Human Genetics, University of Michigan, Ann Arbor, MI

**Keywords:** Genetics, hematology, platelets

## Abstract

SEL1L is a well-known protein in the endoplasmic reticulum associated degradation (ERAD) pathway. While it is known to be expressed in platelets, SEL1L has never been shown to play an active role. Here we find evidence that SEL1L regulates platelet function. We first identified SEL1L through the study of Atypical Equine Thrombasthenia (AET), an autosomal recessive platelet disorder found in Thoroughbred horses. A missense variant in *SEL1L* (c.1810A>G p.Ile604Val) was found in AET-affected horses, which we show is associated with decreased protein expression. SEL1L is intracellular in equine platelets and localizes to the surface upon activation with thrombin. Platelets from homozygous horses exhibit significant decreases in spreading on immobilized collagen. Human megakaryocytes were found to have two SEL1L protein isoforms that increase in expression during megakaryopoiesis, although only one is delivered to mature platelets. Studies using inducible mouse and constitutive zebrafish knockouts demonstrate that SEL1L is necessary for efficient platelet or thrombocyte (fish equivalent) adhesion to sites of endothelial injury. These data reveal a previously undescribed and conserved role for the ERAD pathway in the etiology of AET and platelet function, which may play a role in human platelet disorders as well.

**Brief Summary:** Using a multi-species approach, SEL1L was determined to have a role in platelet function, specifically in helping platelets properly adhere to sites of injury,

## Introduction

Genetic diseases discovered in non-human animals offer the opportunity to discover new genes involved in disease. An inherited platelet dysfunction known as Atypical Equine Thrombasthenia (AET) has been identified in Thoroughbred (TB) horses that is unlike other known genetic bleeding disorders. AET results in reduced platelet response to the physiological agonist thrombin, leading to abnormal bleeding after vascular injury (1,2). Furthermore, this inhibition can result in prolonged bleeding and, in one reported case, epistaxis during racing (1). One prevalence study has been performed and demonstrated that on a single breeding farm one in every 150 TBs was affected (3); however, the frequency of the disease in the general TB population is not known. Pedigrees indicate that the disorder has a recessive mode of inheritance (2).

Coagulation factors have been found to be within normal limits, and platelet counts are normal, suggesting that AET is caused by platelet dysfunction (1). Platelet aggregometry studies, using physiological agonists, demonstrate that AET platelets respond normally to ADP, only respond to collagen with supplementary calcium, and have a minimal response to the most potent agonist, thrombin (1,2). Further research has identified biochemical differences in the AET platelet response to thrombin. After activation, the membrane phosphoinositol pathway is upregulated, resulting in the eventual phosphorylation of protein kinase B (AKT) and release of platelet α-granule contents. AET platelets have disruptions in the phosphoinositol activation pathway, as evidenced by decreased levels of phosphatidylinositol-3,4,5-trisphosphate 5-phosphatase 1 (SHIP1) in the platelet membrane, as well as the increased association of PIK3C2B, a phosphoinositide 3-kinase (PI3K) subunit within the membrane. These two changes likely impact the phosphorylation of AKT and downstream α-granule release (4). α-granules containing factor V (FV) are critical for the acceleration of thrombin activation. AET platelet α-granules contain similar levels of FV to control platelets; however, the amount released is decreased (**Supplemental Fig. 1**) (4). This suggests a failing in the exocytosis of α-granules by activated platelets.

Here we report candidate and genome-wide genetic analysis of AET that identifies a potentially causative missense variant in *SEL1L*. We follow this with functional confirmation in multiple model systems, including horse, human, mouse, and zebrafish. All of these confirm a role of SEL1L in platelet function, which could be the underlying cause of some unknown human bleeding disorders.

## Results

### Platelet fibrinogen binding is defective in AET

Five TBs were identified as AET-affected in previous studies (1–3). They had 16.6%-32.4% fibrinogen bound by platelets as compared to controls (**Fig. 1**). A large degree of intra- and inter-individual variability was identified. All unaffected TBs had >90% fibrinogen bound by platelets compared to controls, except for horse #6, whose dam (horse 3) and full sibling (horse 4) were AET-affected.

**Figure 1:**
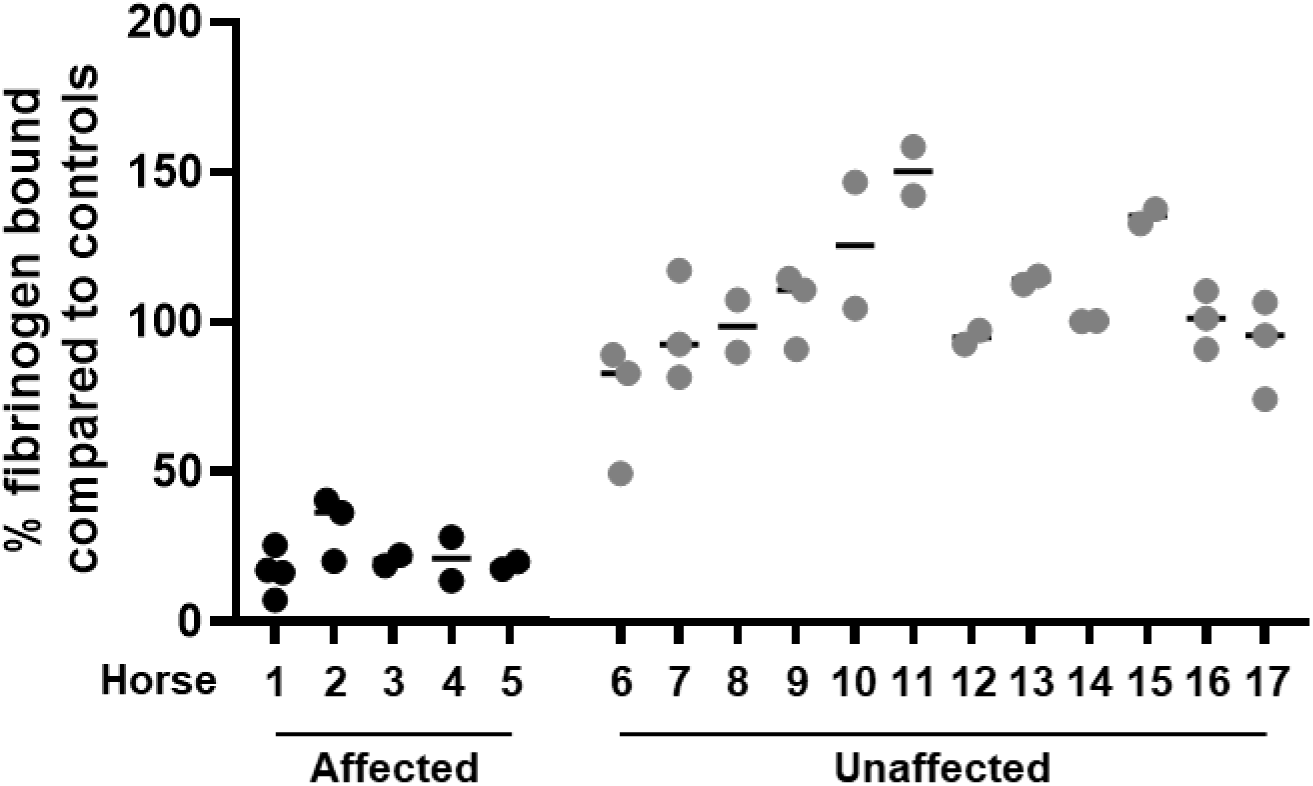
Percent fluorescent fibrinogen bound by thrombin activated platelets for affected (1–5) and unaffected horses (6–17) as compared to the average of three control horses. Each horse was done in duplicate or triplicate. The median for each horse is indicated by the horizontal bar. The affected all have an average percent fibrinogen bound under 35%.

### Candidate gene exclusion

The 5 affected and 12 control horses were whole genome sequenced. All affected horses were confirmed to not have the known equine variants associated with Glanzmann’s Thrombasthenia (5–7). Across the 16 candidate loci, 8,364 variants were identified (**Supplemental Table 2**), but none were significantly associated with AET individuals (*P*>0.05) when either a Bonferroni corrected p-value or a more generous 10% false discovery rate was calculated.

### Whole-genome association identifies variants of interest

Variant callers Freebayes and Delly identified 18,367,554 and 180,965 variants across the genome, respectively. A P<0.0001 cutoff, which allowed half of the controls to be heterozygous, yielded 3,769 Freebayes (**Supplemental Fig. 2**) and 10 Delly variants. Approximately 70% (2,626 variants) of the Freebayes variants were in a ∼6 megabase (Mb) region on chromosome 24 (21Mb - 27Mb; **Supplemental Fig. 2**). All 10 of the Delly variants were in this same region.

Five Freebayes variants were predicted to have a moderate effect on protein function. Only one was in a gene known to be expressed in platelets and interact with other platelet proteins (8) (**Supplemental Table 3**). This was a missense variant in exon 18 of the *suppressor/enhancer of lin-12-like* gene (*SEL1L* c.1810A>G p.Ile604Val*; P*=7.63x10^-6^). Two Delly variants were in or near a gene expressed in platelets. One was a deletion in an uncharacterized lncRNA upstream of *SEL1L,* according to TransMap (genome.ucsc.edu; *AL355838.1:g.26447375_26448962del*; *P*=7.63x10^-6^), and the other was an intronic deletion in *VIPAS39* (g.22685398-22685470del; *P*=3.74x10^-5^).

### RT-PCR confirms gene expression in horse platelets

To confirm that *SEL1L*, the upstream lncRNA, and *VIPAS39* were expressed in equine platelets, end-point PCR was performed with cDNA from unaffected platelets and testicular tissue as a positive control. The observed amplicons demonstrated that *SEL1L* and *VIPAS39* are expressed in equine platelets, but not the upstream lncRNA (**Supplemental Fig. 3**).

### Population study confirms association

A larger population of 114 TBs, including some of the sequenced horses, that had previously been phenotyped (2,3) were genotyped for the *SEL1L* and *VIPAS39* variants. All affected horses were homozygous for the *SEL1L* variant (i.e. alternate allele; Alt/Alt). Platelet binding to fibrinogen was similar between the heterozygous (Ref/Alt) and homozygous WT (Ref/Ref) horses but was significantly different from the Alt/Alt horses (**Fig. 2A**; *P*=2.0x10^-9^). For the *VIPAS39* variant, there was a significant difference in fibrinogen binding between the Alt/Alt horses and the other two genotypes (**Fig. 2B**; *P*=2.0x10^-9^), but one horse below the 35% cutoff was heterozygous.

**Figure 2.**
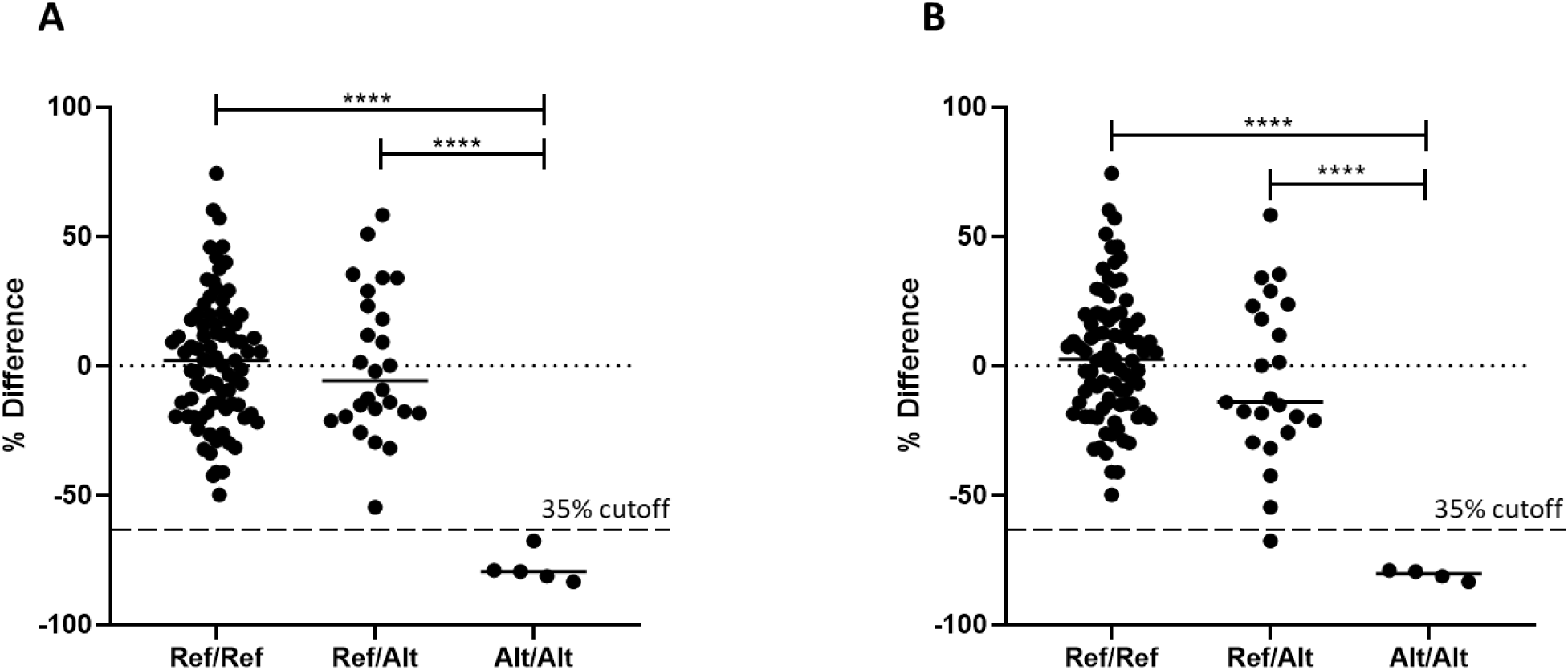
Genotypes of the larger population of 114 phenotyped TBs for (A) *SEL1L* c.1810A>G p.Ile604Val and (B) *VIPAS39:g.22685398_22685470del*. Y-axis is % difference in fibrinogen binding as compared to control horses. The 35% cutoff line characterizing affected horses is indicated by the dashed line. \*\*\*\**P*<0.0001 by ANOVA testing.

### Candidate gene expression in platelets is unaffected by variants

To determine if either variant affected mRNA expression of their respective genes, RT-qPCR was performed on platelet mRNA. Neither *SEL1L* (*P*=0.74) or *VIPAS39* (*P*=0.64) showed a change in expression associated with the variants of interest (**Supplemental Fig. 4**).

Further evaluation of the intronic *VIPAS39* deletion locus in publicly available data from the Functional Annotation of the Animal Genomes project (9) did not reveal any known regulatory elements, suggesting it does not play a role in *VIPAS39* function. Additionally, the AET phenotype is nothing like the Arthrogryposis Renal Dysfunction Cholestasis Syndrome phenotype seen in humans that have known *VIPAS39* deleterious variants (10–12). Thus, the intronic *VIPAS39* deletion was excluded from further study.

### Flow cytometry identifies change in protein expression

Since point mutations can affect protein expression without affecting mRNA levels (13), we investigated SEL1L protein expression using flow cytometry. There were significantly fewer platelets with intracellular SEL1L in horses that were homozygous for the *SEL1L* variant as compared to Ref/Ref (P=0.022; **Fig. 3A**). There also was a non-significant decrease in platelets with intracellular SEL1L between Ref/Ref and Ref/Alt horses (P=0.08). We next questioned whether SEL1L might play a role in platelet interactions following activation. Equine platelets were activated with thrombin, which has been shown to be an aberrant pathway in AET-affected horses (2,4). We found that SEL1L relocated to the platelet surface upon activation (**Fig. 3B**) and there was a trend to have fewer platelets expressing surface SEL1L in AET-affected horses compared to Ref/Ref (P=0.09). However, a significant difference was not observed, likely due to the limited sample size.

**Figure 3.**
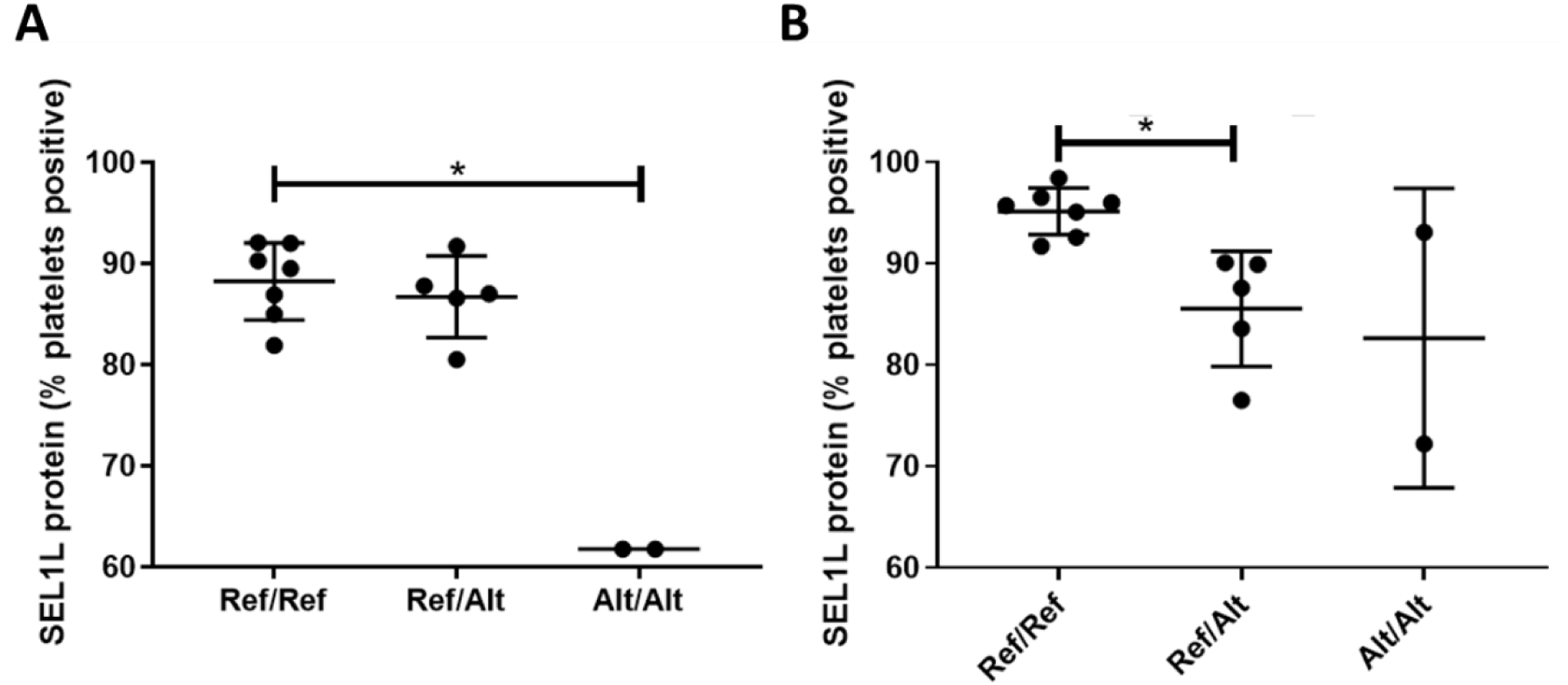
SEL1L protein expression from flow cytometry in resting permeabilized platelets (A) and on the surface of thrombin-activated platelets (B). N = 7 Ref/Ref, 6 Ref/Alt, 2 Alt/Alt. A Kruskal-Wallis test was run with multiple comparisons and a false discovery rate (FDR) correction for multiple testing. *P<0.05

### SEL1L localizes to the surface upon activation and is involved in platelet spreading

We next proceeded to test various coagulation assays and whether there was an association with the *SEL1L* genotypes. This included a template bleeding time (TBT) study, standard clotting assays, and platelet count. No significant difference in TBT between Ref/Ref and Alt/Alt horses (*P*=0.94; **Fig. 4A**) was identified. We also found no differences in plasma clotting assays, fibrinogen levels, or platelet count based on *SEL1L* genotype (**Supplemental Fig. 5**).

**Figure 4.**
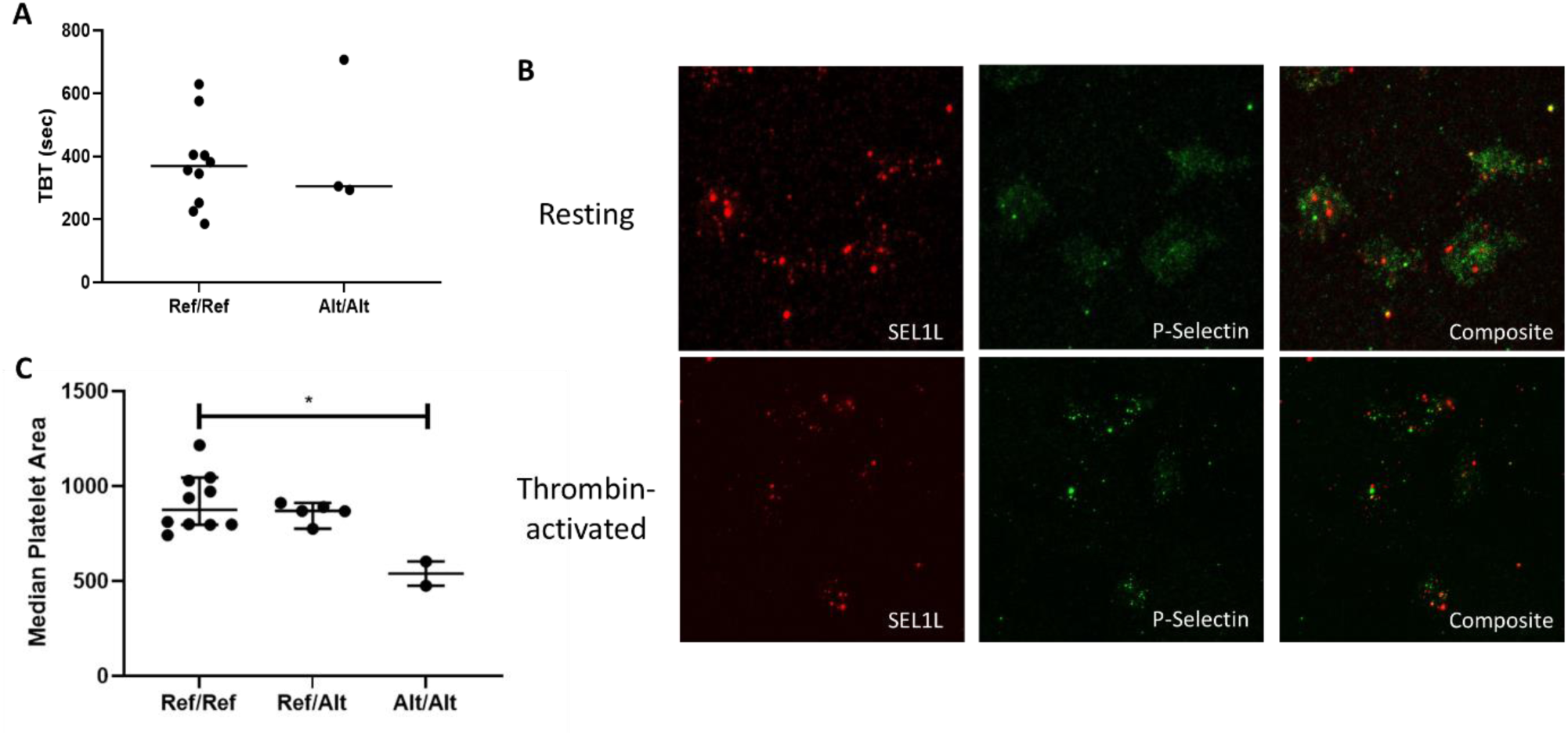
(A) TBTs using horses that were Ref/Ref or Alt/Alt for the *SEL1L* c.1810A>G p.Ile604Val variant found no significant difference between genotypes by Student’s t-test. N= 10 Ref/Ref, 3 Alt/Alt (B) SEL1L (red) localization in relation to P-selectin (green) in (top) resting permeabilized platelets and (bottom) thrombin activated platelet surface of horses that are Ref/Ref for *SEL1L* c.1810A>G p.Ile604Val. SEL1L did not localize to the surface in resting permeabilized platelets (top) but did localize to the surface in thrombin-activated platelets (bottom). N=3 (C) Collagen spreading assay comparing median platelet area to genotype. There was a significant difference between homozygous Ref and Alt genotypes. N=11 Ref/Ref, 5 Ref/Alt, 2 Alt/Alt. A Kruskal-Wallis test with multiple comparisons was run with an FDR correction for multiple testing. \**P<*0.05.

Our flow cytometry data suggest that SEL1L translocates to the platelet surface upon thrombin activation. We therefore evaluated whether it was localized to the α-granule. The membranes of these secretory organelles are incorporated into the platelet membrane during exocytosis (14). Permeabilized resting platelets and thrombin-activated platelets from Ref/Ref and Alt/Alt TBs were fluorescently stained for p-selectin, a marker of α-granules, and SEL1L. The fluorescent markers of these two proteins did not overlap consistently, indicating that SEL1L does not localize to the α-granule in either WT or homozygous mutant platelets (**Fig. 4B; Supplemental Fig. 6**). However, we saw evidence that in both genotypes, SEL1L still localizes to the surface upon thrombin activation.

Lastly, a collagen spreading assay was used to investigate the role of SEL1L in platelet function. First, we confirmed that there was no difference in mean platelet volume between genotypes (*P*=0.45; **Supplemental Fig. 7**). The average area the activated platelets spread was then determined for each horse. A significant decrease in median platelet area was found between Ref/Ref and Alt/Alt horses (*P=*0.023; **Fig. 4C)**.

### SEL1L is expressed by human megakaryocytes and delivered to platelets

To begin to elucidate the mechanism of action of SEL1L in platelets, differentiating human megakaryocytes were used. During megakaryopoiesis, proplatelet forming-megakaryocytes develop nascent platelets at their terminal ends, while boosting gene expression to produce proteins to be delivered to platelets (15).

Proplatelet forming-megakaryocytes were derived from human CD34^+^ hematopoietic stem and progenitor cells (16). During differentiation of cord blood, a progressive increase in the number of cells expressing lineage-specific markers was observed (**Fig. 5A**), paralleled by cell enlargement (**Fig. 5Bi**) and elongation of branched proplatelets (**Fig. 5Bii**). Spontaneous platelet formation from megakaryocytes occurred at day 13 of differentiation when mature platelets are assembled at the terminal ends of the proplatelet branches. Parallel analysis of cell lysates of these megakaryocytes at different days of differentiation showed a significant increase in the expression of SEL1L during megakaryopoiesis, consisting of two variants: SEL1L-p100kDa and SEL1L-p38kDa (**Fig. 5C-D**). Comparable results were obtained in megakaryocytes from human adult peripheral blood progenitors (**data not shown**), indicating that SEL1L expression is present and regulated during megakaryopoiesis, regardless of the source of the hematopoietic progenitors.

**Figure 5.**
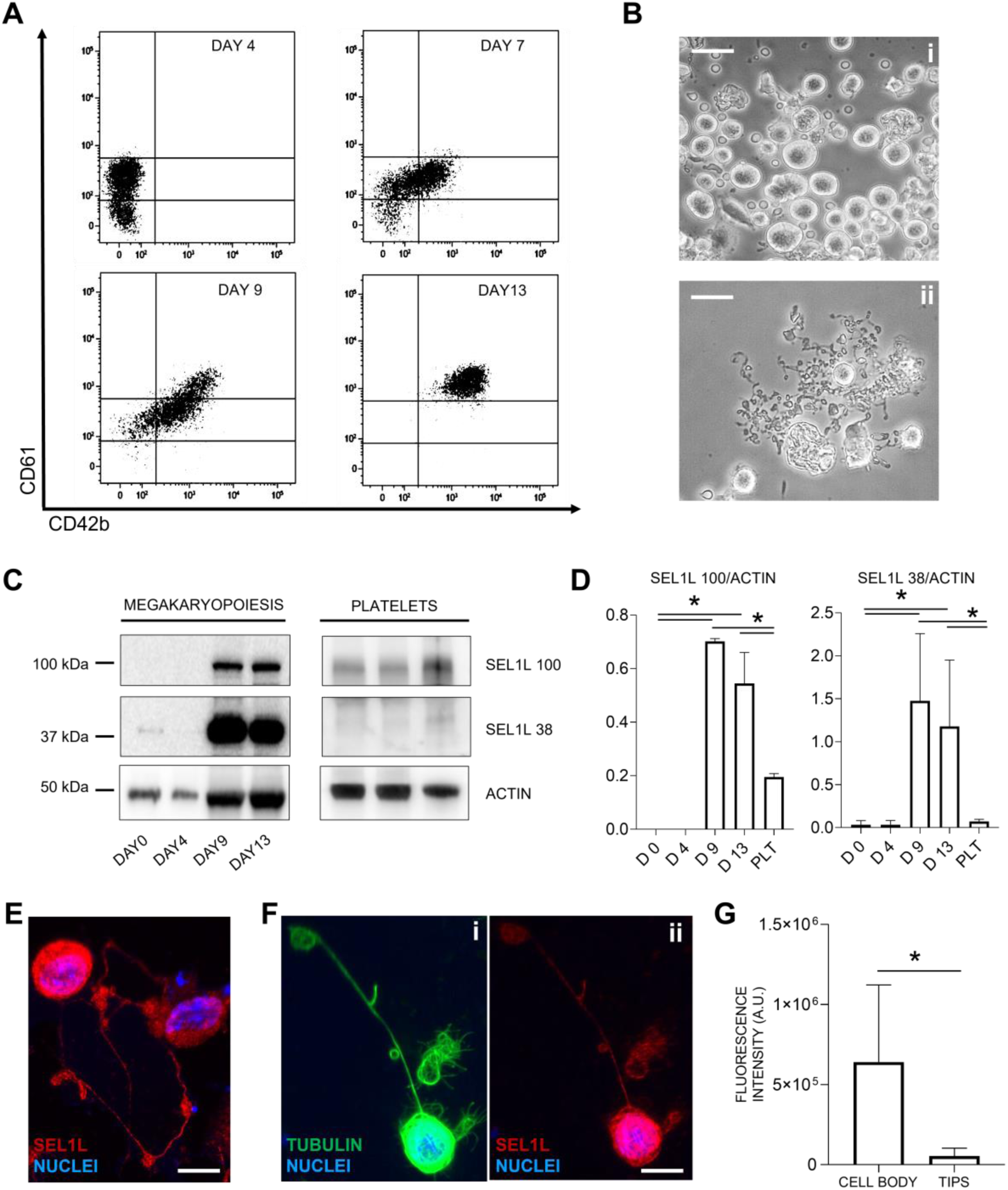
SEL1L is expressed in human megakaryocytes. (**A**) Flow cytometry analyses of human megakaryocytic surface markers from differentiated cord blood. CD61^high^CD42b^high^ megakaryocytes are observed after 2 weeks of differentiation. (**B**) Phase contrast microscopy of mature human megakaryocytes (i) and proplatelets (ii) (scale bar = 50 µM). (**C**) Representative western blot analysis of SEL1L during megakaryopoiesis and in platelets from peripheral blood. (**D**) Densitometric analyses of SEL1L isoforms (n=5, results are presented as mean ± SD). (**E-F**) Immunofluorescence microscopy of megakaryocytes extending proplatelets (scale bar = 30 µM). (**G**) Analysis of SEL1L fluorescence intensity in the megakaryocytic cell body and proplatelet tips. *p<0.05 by Student’s t-test.

While the SEL1L-p100kDa variant remains present in peripheral blood-derived platelets, the level of SEL1L-38kDa decreased to undetectable levels, suggesting that the isoforms are differentially delivered, and mature platelets have different requirements compared to their progenitors (**Fig. 5C-D**). Accordingly, immunofluorescence staining of SEL1L in proplatelet-forming megakaryocytes demonstrated that the protein is mainly localized in the cell body rather than the proplatelet tips (**Fig. 5E-G**). Our data demonstrate that SEL1L expression is prominent during megakaryocyte differentiation, but only one isoform appears to be distributed to mature platelets.

### Loss of SEL1L affects platelet adhesion secondary to endothelial injury *in vivo*

To test whether loss of SEL1L function impacts platelet function, we examined loss of function mutations in mouse and zebrafish models. *Mx1-cre;Sel1L^fl/fl^*mice were used to conditionally knockout the gene in hematopoietic tissue and downstream differentiated cells, followed by examination of platelet binding to collagen. Perfusion of fluorescently-labeled blood over collagen demonstrated a significant loss of platelet adhesion in the *SEL1L^fl/fl^ Mx1-cre^+^* (cre^+^) mice versus control (cre^-^) littermates (*P=*0.03; **Fig. 6**).

**Figure 6.**
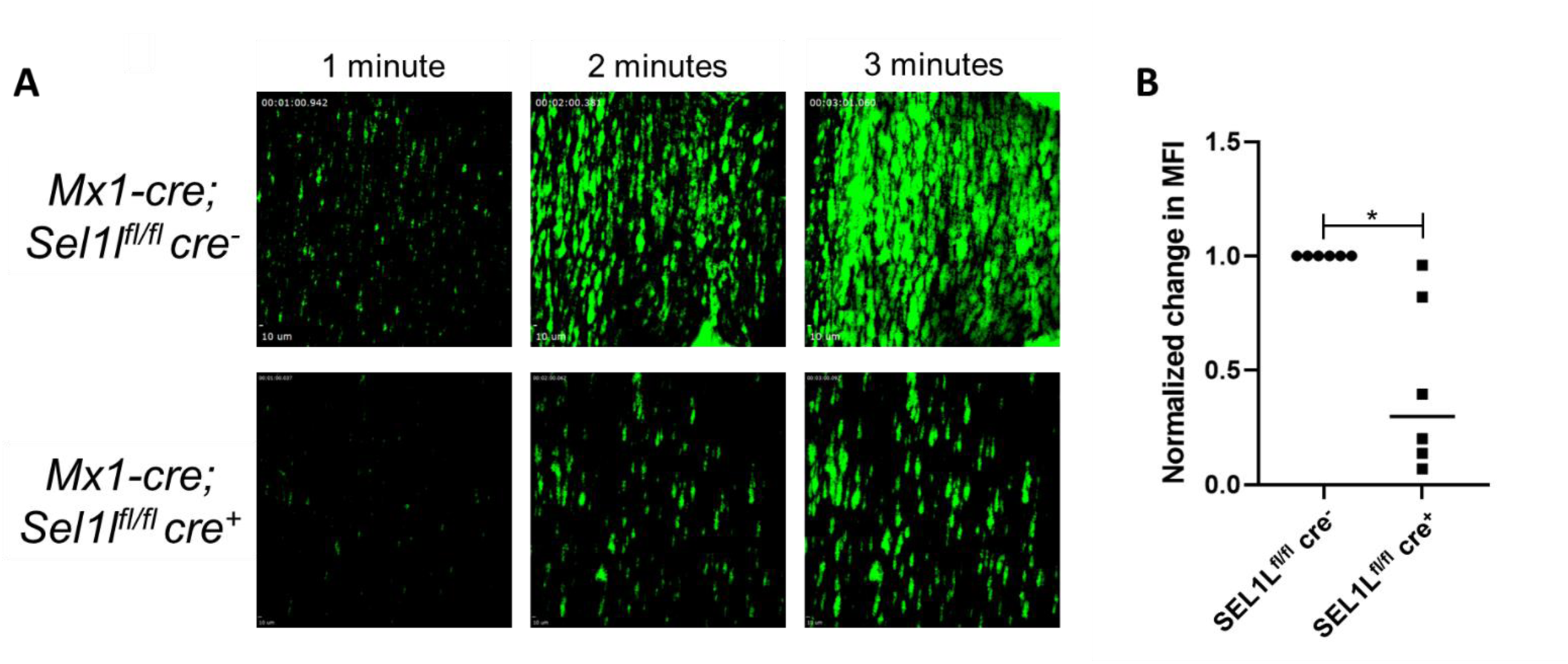
(A) Representative images of murine platelets (green) adhering to a collagen-coated chamber after perfusion of whole blood from both a control SEL1L^fl/fl^ Mx-1 Cre^-^ (cre^-^) and a knockdown SEL1L^fl/fl^ Mx-1 Cre^+^ (cre^+^) mouse. (B) Quantification of the change in mean florescence intensity (MFI) between the beginning and end of the whole blood perfusion across collagen was normalized to the control for each pair. N=6 cre^+^/cre^-^ sibling pairs. A Wilcoxon matched-pairs signed-rank test was performed and found a significant decrease in MFI change between control and knockdown groups, \**P<*0.05.

We next produced a large deletion in zebrafish *sel1l* using CRISPR/Cas9 genome editing, deleting nearly the entire coding region (**Supplemental Fig. 8**). Laser-mediated arterial endothelial injury was performed on 5 days post fertilization (dpf) offspring of a heterozygous incross. While there was no significant difference in time to occlusion or time to first thrombocyte attachment, there was a significant difference in total number of thrombocytes that adhere (*P=*0.02; **Fig. 7**). There were no significant differences in total thrombocyte counts (**Supplemental Fig. 9**).

**Figure 7.**
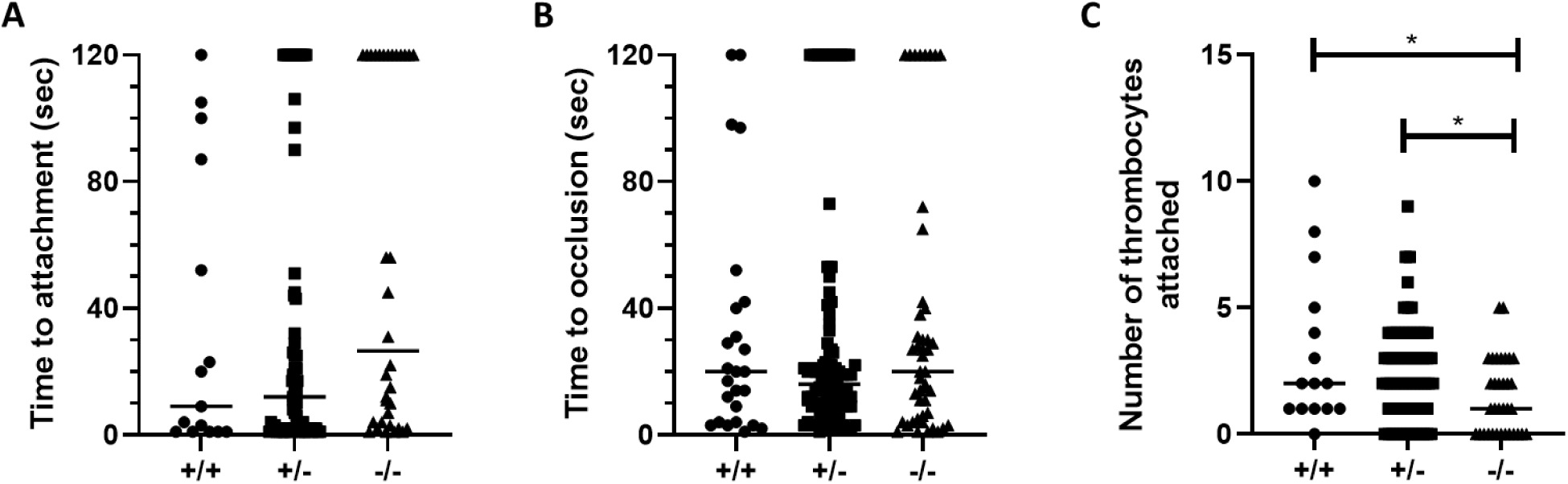
Laser-mediated arterial endothelial injury of a heterozygous incross at 5 dpf shows no difference (A) in time to first thrombocyte attachment (N= 15 +/+, 69 +/-, 34 -/-.) or (B) time to occlusion between wild-type (+/+), heterozygous (+/-), or homozygous (-/-) larvae (N= 25 +/+, 89 +/-, 46 -/-.), but there was a (C) significant decrease in the number of thrombocytes that attach to the site of injury in homozygous larvae as compared to those that were wild-type and heterozygous (N= 15 +/+, 69 +/-, 34 -/-). \**P<*0.05 by Kruskal-Wallis testing with multiple comparisons and an FDR correction.

## Discussion

This study initially investigated the genetic etiology of AET and followed up by studying the previously unknown role of SEL1L in platelet function. AET is an inherited platelet disorder that prevents affected horses from clotting efficiently following vascular injury. Whole-genome sequencing identified a missense variant in *SEL1L (*c.1810A>G p.Ile604Val). This variant was associated with expression changes at the protein level and decreased platelet spreading on collagen. Human megakaryocyte studies determined protein localization and identified two isoforms. Finally, mouse and zebrafish *SEL1L* knockout models definitively identified a functional role for SEL1L in platelet function. Therefore, the missense variant is likely the cause of AET.

A limitation throughout this study was the number of horses available for study. We only identified *n*=3 TBs that were Alt/Alt for the *SEL1L* c.1810A>G p.Ile604Val variant and that were readily accessible, since fresh platelets or horses were required for most of the equine experiments. The TBT has a high degree of both intra-and inter-horse variability (17) which likely led to the non-significant results. Additionally, the high TBT seen in the index case (>120 minutes) may have been primarily due to the consumption of coagulation factors as it was evaluated after bleeding for three weeks (1). We also demonstrated that, while SEL1L is an intracellular protein, it does not localize to the α-granule in resting platelets, but it does relocate to the plasma membrane upon thrombin stimulation. This indicates that SEL1L is likely localized to an organelle within the platelet that is involved in exocytosis post-activation, such as dense or lysosomal granules. Since SEL1L is known to localize to the endoplasmic reticulum in nucleated cells (18), we hypothesize that platelet SEL1L is located in the dense tubular system (DTS), a remanent of the progenitor megakaryocyte endoplasmic reticulum (ER) (19,20). The DTS contains proteins known to be involved in protein folding such as protein disulfide isomerase (21), and at least one of these has been shown to move to the platelet surface upon activation (22). SEL1L may be migrating to the cell membrane upon activation in a similar manner.

Three additional model systems were used to further elucidate the role of SEL1L in platelets. First, SEL1L was investigated in human megakaryocyte cell culture. SEL1L was found to be expressed in two isoforms that increase in expression during megakaryopoiesis. Both isoforms showed a significant increase in expression during megakaryopoiesis, with only one isoform, SEL1L-p100kDa, appearing to be passed on to mature platelets. The distribution of proteins from megakaryocytes to platelets is tightly regulated. Some of the key regulators of megakaryopoiesis and proplatelet formation, including ATXN2, P2Y13, and uridine diphosphate (UDP)-galactose-4-epimerase (GALE), have been shown to be primarily expressed in megakaryocyte progenitors rather than mature platelets (23–25). As such, while two SEL1L isoforms are expressed in megakaryocytes, the transfer of the larger isoform to platelets suggests it may play the functional role in mature platelets.

SEL1L was also investigated in mouse and zebrafish models, demonstrating that it is necessary for proper platelet/thrombocyte adhesion to an injured vessel. Since the *SEL1L* knockout is embryonic lethal in mice (26,27), we took advantage of a conditional mutant. However, we found that zebrafish with complete loss of SEL1L survive into adulthood. Previous work in zebrafish using morpholinos found that *sel1l* knockdown led to cardiac and brain edema, decreased blood circulation, and blood stasis in the tail (28), which had suggested that SEL1L does play a role in circulation. Our data (not shown) suggest the possibility that these morpholino phenotypes could be the result of off-target effects, or genetic compensation allowing rescue in the knockout (29).

We found that SEL1L plays a role in platelet activation and adhesion. In other cell types, it has been shown that the primary function of SEL1L is in ER-associated degradation (ERAD) (26,27). SEL1L forms a complex with Hrd1 (30) and is necessary for ER homeostasis and proper protein degradation and secretion (26,27). A parallel and cooperative pathway to ERAD is the unfolded protein response (UPR) (31–33). UPR activation has been identified in mammalian platelets (34), thus, SEL1L could play a similar role in protein degradation in platelets. Recent studies have identified novel mechanisms of SEL1L modulating cell-signaling pathways by regulating the ubiquitination of a transcription factor and controlling the degradation of a specific receptor (35,36). SEL1L may have a similar mechanism in AET, where it regulates a protein involved in platelet function, likely in the thrombin signaling pathway since that is where differences were previously identified in AET affected horses (2,4).

In addition to SEL1L’s established role in ERAD, two other truncated isoforms of SEL1L have been shown to not solely localize to the ER in human lymphocytes (37). The largest isoform was predicted to be ∼89kDa, and the truncated ones were 34 and 35kDa. Similarly, we identified two isoforms of SEL1L in megakaryocytes, 100 and 38kDa. The 100kDa isoform is transferred from megakaryocytes to platelets at low levels, and further study is necessary to determine if this isoform functions outside of the traditional ERAD pathway.

In summary, these results suggest that *SEL1L* c.1810A>G p.Ile604Val is responsible for the etiology of AET. SEL1L appears to be necessary for platelets or thrombocytes to properly adhere to sites of injury through interaction with collagen, and loss of function impacts hemostasis. This identification of the putatively causative variant of AET will allow for affected TBs and carriers to be identified and inform breeding decisions. Additionally, this work has identified SEL1L, and possibly the ERAD pathway, as important players in platelet function, which could potentially have implications for patients with unknown bleeding disorders.

## Methods

### Sex as a biological variable

Our experiments used both male and female animals as there was no difference in phenotype between sexes.

#### Horses and whole genome sequencing

Seventeen TBs (5 affected, 12 unaffected) were phenotyped in a previously published study (3) Briefly, platelets were activated with thrombin, the amount of fibrinogen bound was quantified via flow cytometry and then compared to the average of three unaffected horses (2). A strict cut-off of an average of 35% of fibrinogen bound by platelets or less compared to control horses was used to definitively diagnose affected horses. Whole-genome sequencing was performed on the entire group using an Illumina HiSeq 2500, at a targeted 50x coverage with paired end 150 base-pair (bp) reads. Raw reads were trimmed with Trimmomatic (38), mapped to the EquCab3.0 reference genome (39) using Burrows-Wheeler Aligner (BWA-MEM) (40), and sorted by coordinate with samtools (41). PCR duplicates were removed with Picard tools (http://broadinstitute.github.io/picard/). Integrative Genome Viewer (IGV) (42), was used to genotype all horses for the two variants shown to cause Glanzmann’s Thrombasthenia (5–7). Freebayes (43) was used to call single nucleotide variants (SNVs) and small insertions/deletions. Delly was used to identify larger structural variants (44). The functional effects of the variants were predicted using SnpEff (45), and the variants were filtered by segregation using Fischer’s Exact Test with SnpSift (46). Associated variants were identified and filtered by predicted effect. PlateletWeb (http://plateletweb.bioapps.biozentrum.uni-wuerzburg.de/plateletweb.php) was used to identify the variants that are in and near genes known to be expressed in human platelets. KaryoploteR (47) was used to identify haploblocks among the variants that were *P*<0.0001.

#### Candidate gene approach

Candidate genes (*AKT1, AKT2, F5, SHIP1, PIK3C2B, F2, F2R, PIK3C2A, PIK3C2G, PIK3CA, PIK3CB, PIK3CG, PIK3R1, PIK3R4, PIK3R5, PIK3R6*) were identified from previous biochemical work (4) on AET-affected platelets (**Supplemental Fig. 1**). One kilobase of sequence up and downstream was included to account for the 5’ and 3’ untranslated regions that could harbor regulatory variants. The functional effects of the variants were predicted using SnpEff (45), and variants were filtered by segregation using Fisher’s Exact Test with SnpSift (46).

#### Equine blood processing

Blood was drawn into acid citrate dextrose tubes and platelets were separated from whole blood by density centrifugation to yield washed platelets. To remove leukocytes, the washed platelets were treated with prostaglandin E1, and leukocyte depleted using an anti-equine F6 antibody generously provided by Dr. Jeff Stott.

#### RNA isolation, reverse transcription PCR, and quantitative PCR (RT-qPCR)

RNA was extracted from leukocyte-depleted platelets using phenol-chloroform phase separation, as previously described (48) and treated with Turbo DNase (Thermo Fisher Scientific, Waltham, MA). Superscript III (Thermo Fisher Scientific, Waltham, MA) was then used to reverse transcribe 500 ng of RNA. Primers that yield products spanning two exons were designed for *SEL1L, AL355838.1*, and *VIPAS39* (**Supplemental Table 1**) and end-point PCR was performed on a sample from a wild-type (WT) horse.

Complementary DNA from leukocyte-depleted platelets (*SEL1L A>G*: *n*=12 WT, 6 heterozygous, 2 homozygous; *VIPAS39del n*=16 WT, 3 heterozygous, 1 homozygous) was run in triplicate using SYBR green on an AriaMx Real-Time PCR System (Agilent, Santa Clara, CA). The gene *ACTB* was used for normalization (**Supplemental Table 1**), and ΔΔCt values were calculated.

#### Genotyping additional horses

114 phenotyped (3) TBs were genotyped for *SEL1L* A>G using polymerase chain reaction-restriction fragment length polymorphism (PCR-RFLP) with the restriction enzyme HpyCH4III or for the *VIPAS39* deletion by visualization on an agarose gel (**Supplemental Table 1**).

#### Flow cytometry of equine platelets

Washed equine platelets (*SEL1L A>G*: *n*=7 WT, 5 heterozygous, 2 homozygous alternate) were fixed in 1% paraformaldehyde, permeabilized in 0.1% NP-40 and labeled with a monoclonal SEL1L antibody conjugated to AlexaFluor488 (Santa Cruz Biotechnology, Dallas TX) and a monoclonal P-Selectin antibody conjugated to APC (Invitrogen, Waltham, MA). Samples were analyzed on a Beckkman Coulter FC500 flow cytometer and data analyzed with FlowJo.

#### Template bleeding times

Template bleeding times (TBTs) were measured on 13 horses (10 WT, 3 homozygous alternate) as previously described (49). Briefly, a template system (MedexSupply, Passaic NJ) was used on a clipped area just distal to the accessory carpal bone on the caudomedial and caudolateral aspects of the forelimbs. A sphygmomanometer cuff was placed proximal to the carpus and inflated to 40 mmHg for 60 seconds to achieve cutaneous venostasis. Blood flow from the incision was collected on a No. 40 absorbent filter paper at 30 second intervals until no capillary flow was absorbed. Bleeding times were performed at a standardized time each morning on both forelimbs over three consecutive days and mean times determined. Blood samples for complete blood count and basic coagulation panel (prothrombin time, activated partial thromboplastin time, and fibrinogen) were obtained prior to the study to ensure no influences by changes in platelet count or clotting defects.

#### Immunofluorescence

Resting and thrombin activated (bovine alpha-thrombin, Haematologic technologies, Essex Junction, VT) washed TB platelets (10 x 10^6^ plt/mL) were incubated on poly L-lysine coverslips (37°C/1hr). The platelets were then fixed with 1% paraformaldehyde, permeabilized with 0.1% NP-40, and blocked with 10% bovine serum albumin. Platelets were incubated with anti-P-selectin (BD Pharmaceuticals, diluted 1:10 in 5% goat serum), followed by a secondary antibody conjugated to an AF488 fluorophore (Thermofisher, Waltham, MA; diluted 1:20 in 5% goat serum). Both incubations were performed for 2 hours rocking at room temperature. Next, fixed platelets were incubated with a cocktail of anti-SEL1L and anti-CD42b (Abcam, Waltham, MA; diluted 1:200 and 1:10 respectively in 5% goat serum) rocking overnight at 4°C. Lastly, the platelets were incubated with a cocktail of secondary antibodies conjugated to AF555 and AF405 (Thermofisher, diluted 1:300 and 1:10 respectively in 5% goat serum) for 2 hours rocking at room temperature. The stained platelets were imaged on a Leica TCS SP8 STED 3X (Leica Microsystems, Wetzlar, Germany).

#### Collagen spreading assay

Washed platelets (9 x 10^9^ plt/L) were allowed to adhere to type 1 collagen coated coverslips (Electron Microscopy Sciences, Hatfield PA; 37°C/2hrs), washed in buffer, and fixed in 1% paraformaldehyde (room temperature/1hr). The mean platelet volume of the platelet-rich plasma (*n*=11 WT, 5 heterozygous, 2 homozygous alternate) was determined. The platelets were washed and allowed to adhere to type 1 collagen coated coverslips (9 x 10^9^ plt/L; Electron Microscopy Sciences, Hatfield PA; 37°C/2hrs). These were subsequently washed in buffer, fixed in 1% paraformaldehyde (room temperature[RT]/1hr), and mounted. The platelets were imaged using an Olympus BX62 microscope. Platelet area was determined using ImageJ 1.51K (http://imagej.nih.gov/ij) by an observer blinded to genotype.

#### Human megakaryocyte differentiation

Umbilical cord blood units were collected in bags containing acid-citrate-dextrose (ACD). Megakaryocytes were differentiated from human CD34^+^ cells using previously described methods (16). Briefly, CD34^+^ cells were separated by an immunomagnetic sorting technique (50). The purity of the sorted population was routinely assessed by flow cytometry to be 90±4% of the total cells recovered. CD34^+^ cells were cultured for 13 days in Stem Span media (STEMCELL Technologies, Vancouver, Canada) supplemented with 10 ng/mL TPO and interleukin-11 (Peprotech, London, UK); 1% penicillin-streptomycin and 1% L-glutamine (Euroclone, Milan, Italy). The culture medium was renewed every three days.

#### Isolation of human platelets from peripheral blood

Human whole blood from healthy subjects was collected using ACD and processed as previously described (51). Briefly, platelets were isolated by centrifugation at 200g for 10 minutes to obtain platelet-rich plasma (PRP). PRP was then centrifuged at 1500g for 15 minutes in presence of 0.2 U/mL apyrase grade I and 1 µm PGE1 (Sigma-Aldrich). Platelets were then gently resuspended in Tyrode’s buffer (134 mM NaCl; 0.34 mM Na2HPO4; 2.9 mM KCl; 12 mM NaHCO3; 20 mM HEPES; 5 mM glucose; pH 6.5). Washed platelets were finally lysed with Hepes-glycerol lysis buffer (Hepes 50 mM, NaCl 150 mM, 10% glycerol, 1% Triton X-100, MgCl2 1.5 mM, EGTA 1 mM, NaF 10 mM, PMSF 1 mM, Na3VO4 1 mM, 1 µg/ml leupeptin, 1 µg/ml aprotinin). Before being lysed, purity of the platelet samples was assessed by flow cytometry, as previously described (52).

#### Immunoblotting

Human platelet and megakaryocyte protein lysates were subjected to gel electrophoresis (SDS-PAGE) and transferred to polyvinylidene fluoride (PVDF) membrane (BioRad, Milan, Italy). Membranes were probed with mAb anti-SEL1L (53) (1:200) and mAb anti-βactin (1:1000). Immunoreactive bands were detected by horseradish peroxidase-labeled secondary antibodies (BioRad), using enhanced chemiluminescence reagent (Millipore). Pre-stained protein ladders (BioRad) were used to estimate the molecular weights. Band intensities from individual western blots were quantified by densitometric analysis using the ImageJ.

#### Analysis of proplatelet formation

Megakaryocyte and proplatelet yields were evaluated at the end of the cell culture as previously described (16). Briefly, 1x10^5^ cells were seeded in a 24-well plate and incubated at 37°C in a 5% CO2 fully humidified atmosphere. Proplatelets were identified as cells displaying long filamentous structures, ending with platelet-sized tips. Phase contrast images were obtained by an Olympus IX53 microscope (Olympus).

For the immunofluorescence analysis, mature megakaryocytes were seeded onto 12 mm glass coverslips coated with 25 µg/mL human fibronectin (Corning Inc., Bedford, MA, USA) for 2 hours at room temperature (RT) and subsequently blocked with 1% albumin (Sigma Aldrich, Milan Italia) for 1 hour at RT. Megakaryocytes were harvested and plated onto coated coverslips in 24-well plates and allowed to adhere for 16 hours at 37**°**C in a 5% CO2 fully humidified atmosphere. Next, cells were fixed in 4% paraformaldehyde (Sigma Aldrich, Milan, Italy), permeabilized with 0.5% Triton X-100 (Sigma Aldrich, Milan, Italia), blocked with 5% BSA and stained with anti-β1-tubulin antibodies (1:1000) and pAb SEL1L (1:200) for 1 hour at RT, and the Alexa Fluor secondary antibodies (1:500, Invitrogen, Milan, Italy) for 2 hours at RT. Nuclear counterstaining was performed using Hoechst 33258 (100 ng/ml, Sigma Aldrich, Milan, Italy) for 3 minutes at RT. Specimens were mounted in ProLong Gold Antifade Reagent (Invitrogen, Milan, Italia). Negative controls were routinely performed by omitting the primary antibody. Immunofluorescence images were acquired by a Nikon Ti2-JU. Megakaryocytes extending proplatelets were identified as cells extending tubulin-positive long filamentous structures ending with platelet-sized tips.

#### Flow cytometry analysis of megakaryocytes

For megakaryocyte differentiation analysis, cells were suspended in PBS buffer and stained with antibodies with FITC anti-CD61 and PE anti-CD42b antibodies, at RT, in the dark for 30 minutes. All samples were acquired with a BD flow cytometer. Off-line data analysis was performed using Kaluza software package.

#### Murine SEL1L conditional knockdown and platelet perfusion over collagen

*Mx1-cre^+^;Sel1L^fl/fl^* and *Mx1-cre^-^;Sel1L^fl/fl^*mice were treated at 10 weeks with poly-inosine-poly-cytosine (pIpC) as previously described (54). Using cre^+^ and cre^-^ paired littermates, whole blood was collected from the retro-orbital space into sodium citrate and incubated with DiOC6 (ThermoFisher Scientific). Platelet adhesion was measured as previously described (55,56). Briefly, flow chamber μ-slides (1 × 17 mm; Ibidi) were coated with 50 μg/mL equine collagen (Chronolog) overnight at 4°C and rinsed with 37°C 1X phosphate buffered saline. The blood was perfused over the collagen coated chamber at 1,800 s^−1^, a typical arterial shear rate, by an automated syringe pump. Adhesion of the fluorescently labeled platelets was recorded using a Zeiss Axiovert 200M inverted fluorescent microscope at 40x. The videos were analyzed using 3i Slideobook. The mean fluorescence was measured at the beginning and end of perfusion. The difference was determined and normalized to the cre^-^ mouse of each pair.

#### Development of sel1l knockout zebrafish

A *sel1l* knockout zebrafish line was generated on a wild-type background of a hybrid of AB and TL strains (ABxTL). Synthego’s CRISPR Design Tool was used to identify CRISPR/Cas9 single guide RNA (sgRNA) target sites in exons 3 (GAAGGUGCCAAUAUUGUGAC) and 20 (GGAUCUCAAUGUAAGUUGAG). The sgRNA was complexed with Cas9 (Synthego) and injected into one-cell embryos and raised to adulthood. The fish were genotyped and crossed to *cd41-eGFP* transgenic fish (57) to establish a line with fluorescently labeled thrombocytes.

#### Laser-mediated endothelial injury in larval zebrafish

At 5 days post fertilization (dpf) laser-mediated arterial endothelial injury was performed as previously described (58,59). Briefly, the larvae were anesthetized with tricaine and mounted with 0.8% low melting temperature agarose on a glass coverslip. Endothelial injury was performed with a pulsed nitrogen dye laser system (Andor Technology) and analyzed on an inverted microscope (Olympus IX73) using a 20x objective. The endothelium of the dorsal aorta at the 5^th^ somite posterior to the anal pore was targeted with the laser. *cd41-eGFP* + thrombocytes were visible under fluorescence and the time to first thrombocyte attachment (TTA) was measured (up to 2 minutes). The total number of thrombocytes that attached and time to full occlusion (TTO; up to 2 minutes) were also measured. The larvae were genotyped after the data were collected.

#### Thrombocyte counting

Relative thrombocyte count was measured as previously described (60). Briefly, anesthetized 5 dpf larvae were immobilized in 0.8% low melt agarose in a glass capillary. The capillary was submerged in water and placed under a Leica MZ16FA stereo microscope with GFP fluorescence. A one-minute video was recorded and VirtualDub (https://www.virtualdub.org/) used to transform the videos into grayscale, then ImageJ used to analyze the relative thrombocyte count in each video.

### Statistical analysis

Analysis of data was performed using GraphPad Prism software. Parametric data were analyzed with an ANOVA or Student’s t-test. Nonparametric data were analyzed with a Kruskal-Wallis test with multiple comparisons and a false discovery rate (FDR) correction or a Wilcoxon matched-pairs signed-rank test. A statistically significant difference was considered P<0.05.

#### Study approval

All horse experiments were conducted in accordance with guidelines approved by the University of California Davis Animal Care and Use Committee (ACUC). All human megakaryocyte experiments obtained written informed consent at enrollment in accordance with the ethical committee of the I.R.C.C.S. Policlinico San Matteo Foundation of Pavia and the principles of the Declaration of Helsinki. All mouse and zebrafish experiments were done in accordance with guidelines approved by the University of Michigan ACUC.

### Data availability

All sequencing reads for this study have been submitted to NCBI Sequence Read Archive, PRJNA553581. Full VCF files for each candidate gene were submitted to EVA, PRJEB33757.

## Author contributions

A.R.D designed research studies, conducted experiments, acquired data, analyzed data, and wrote manuscript.

F.C conducted experiments, acquired data, and analyzed data.

J.W.N designed research studies, conducted experiments, acquired data, and analyzed data.

R.A designed research studies, conducted experiments, and acquired data.

E.M.K conducted experiments, acquired data, and provided reagents.

Q.L designed research studies and provided reagents.

M.H designed research studies and provided reagents.

I.B Designed research studies, analyzed data, and provided reagents.

A.B Designed research studies, analyzed data, and provided reagents.

J.A.S designed research studies, provided reagents, and edited manuscript.

F.T designed research studies, conducted experiments, acquired data, analyzed data, provided reagents, and edited manuscript.

C.J.F designed research studies, provided reagents, and edited manuscript.

## Acknowledgements

This work was supported by National Institutes of Health grants T32 HL007622 (J.A.S), R35 HL150784 (J.A.S), R01 ES 032255 (J.A.S), 1K01OD015134 (C.J.F). A.R.D was supported by the Ann T. Bowling Fellowship, Center for Equine Health, Louis R. Rowan Fellowship, and Boxer Training Program in Molecular and Translational Hematology. J.A.S is the Henry and Mala Dorfman Family Professor of Pediatric Hematology/Oncology and receives support from the American Society of Hematology and National Hemophilia Foundation.

The authors would like to thank the University of California Center for Equine Health for use of their horses. They would like to thank Franco Marincola and Qian Zhang for DNA sequencing courtesy of the Sidra Medical and Research Center. They would also like to thank Dr. Mike Lenardo for his support as well as Dr. Tatiana Viau, the staff at CEH, Nghi Nguyen, Dr. Callum G. Donnelly, Allison Ferguson, M. Hao Hao Pontius, Ashley Keesling, and Katherine Sowell for their technical support.

## Notes

**Conflict of interest disclosures:** J.A.S. has been a consultant for Sanofi, Takeda, Genentech, CSL Behring, and HEMA Biologics. M.H. is a consultant and equity holder for Veralox Therapeutics and Cereno Scientific. A.R.D., F.C., J.W.N., R.A., E.M.K., Q.L., I.B., A.B., F.T., C.J.F. have no conflicts of interest to declare.

### Competing Interest Statement

J.A.S. has been a consultant for Sanofi, Takeda, Genentech, CSL Behring, and HEMA Biologics. M.H. is a consultant and equity holder for Veralox Therapeutics and Cereno Scientific. A.R.D., F.C., J.W.N., R.A., E.M.K., Q.L., I.B., A.B., F.T., C.J.F. have no conflicts of interest to declare.

