## Supplemental Figures for "Identification of the role of SEL1L in platelet function through a multi-species genetic investigation"

### Supplemental Figure 1

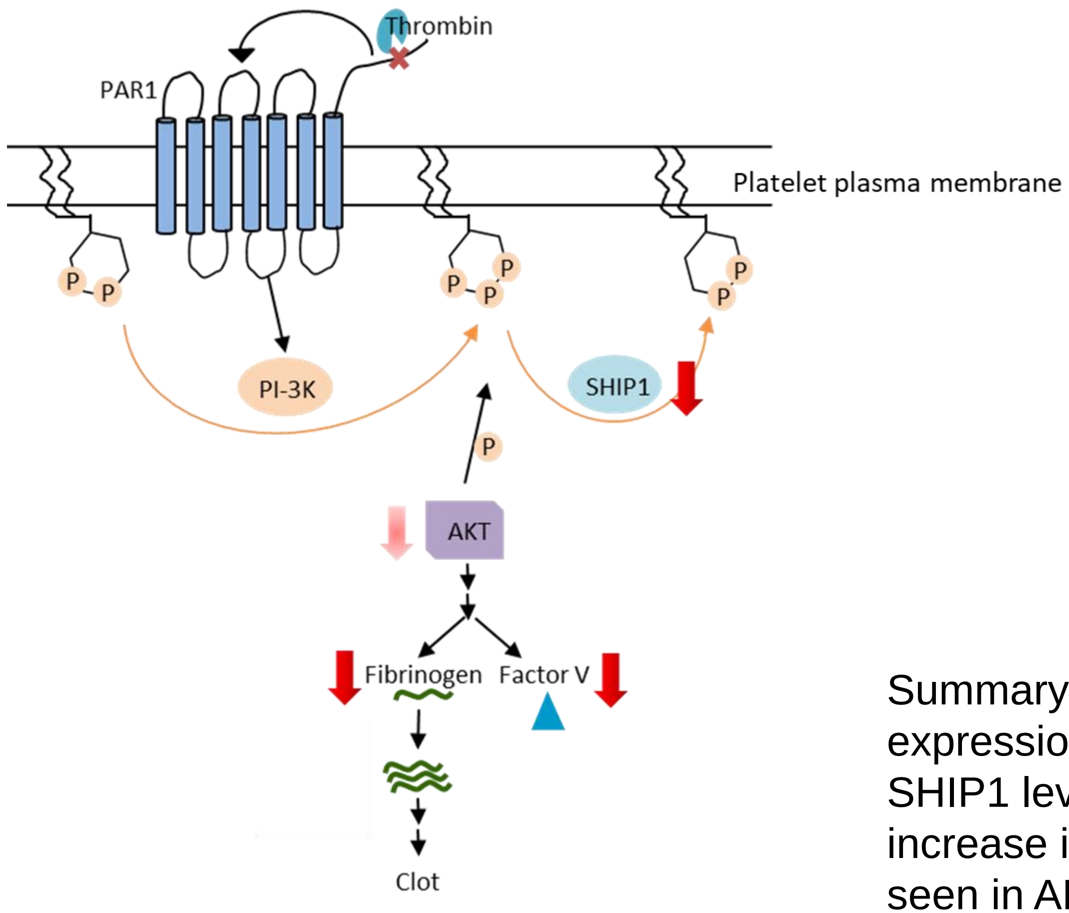


Summary of thrombin signaling pathway. Changes in protein expression in AET-affected horses are indicated by red arrows. SHIP1 levels are decreased in the platelet membrane, and an increase in association of PIK3C2B with the membrane were seen in AET-affected horses. There is abnormal phosphorylation of AKT after activation as well as decreased levels of fibrinogen bound by activated platelets. Additionally, the amount of FV released is decreased in affected horses.

### Supplemental Table 1

| **Table 1.S1.** Primer Sequences | | | |
|---|---|---|---|
| **Primer Target** | **Coordinates** | **Forward** | **Reverse** |
| *SEL1L* transcript | chr24:26,074,490-26,076,942 | CCTAGCACTCTGCAAATTAGGC | CAAGTTGGGTGAAGACATCTCG |
| *VIPAR* transcript | chr24:22,684,454-22,686,432 | CAGGACCCTGAGAAACGAAA | CCAGGAGCGTGTAATGGTCT |
| lncRNA transcript | chr24:26,327,936-26,328,024 | CTGCTGAGCTCTCTGGGTGT | CAGCTGGAGTGACTGAGCATT |
| *SEL1L* genotyping | chr24:26,080,981-26,082,166 | CAACGTCAGTGCCAAATCC | ACATTGCCTTTCCAGCAGTC |
| *VIPAR* genotyping | chr24:22,685,271-22,685,559 | TGAGCCACAGCCACTTGTTA | TTCCAGACCCGTCTAGCATC |
| *ACTB* transcript | chr13:4,382,919-4,383,011 | AAGGAGAAGCTCTGCTATGTCG | GGGCAGCTCGTAGCTCTTC |

### Supplemental Table 2

| Number of variants identified in candidate genes | |
|---|---|
| Gene | Total variants |
| *AKT1* | 251 |
| *AKT2* | 181 |
| *F2* | 166 |
| *F2R* | 64 |
| *F5* | 448 |
| *FGA, FGB* | 152 |
| *PIK3C2A* | 329 |
| *PIK3C2B* | 421 |
| *PIK3C2G* | 2598 |
| *PIK3CA* | 97 |
| *PIK3CB* | 688 |
| *PIK3CG* | 128 |
| *PIK3R1* | 374 |
| *PIK3R4* | 1144 |
| *PIK3R5, PIK3R6* | 602 |
| *SHIP1* | 721 |

### Supplemental Figure 2

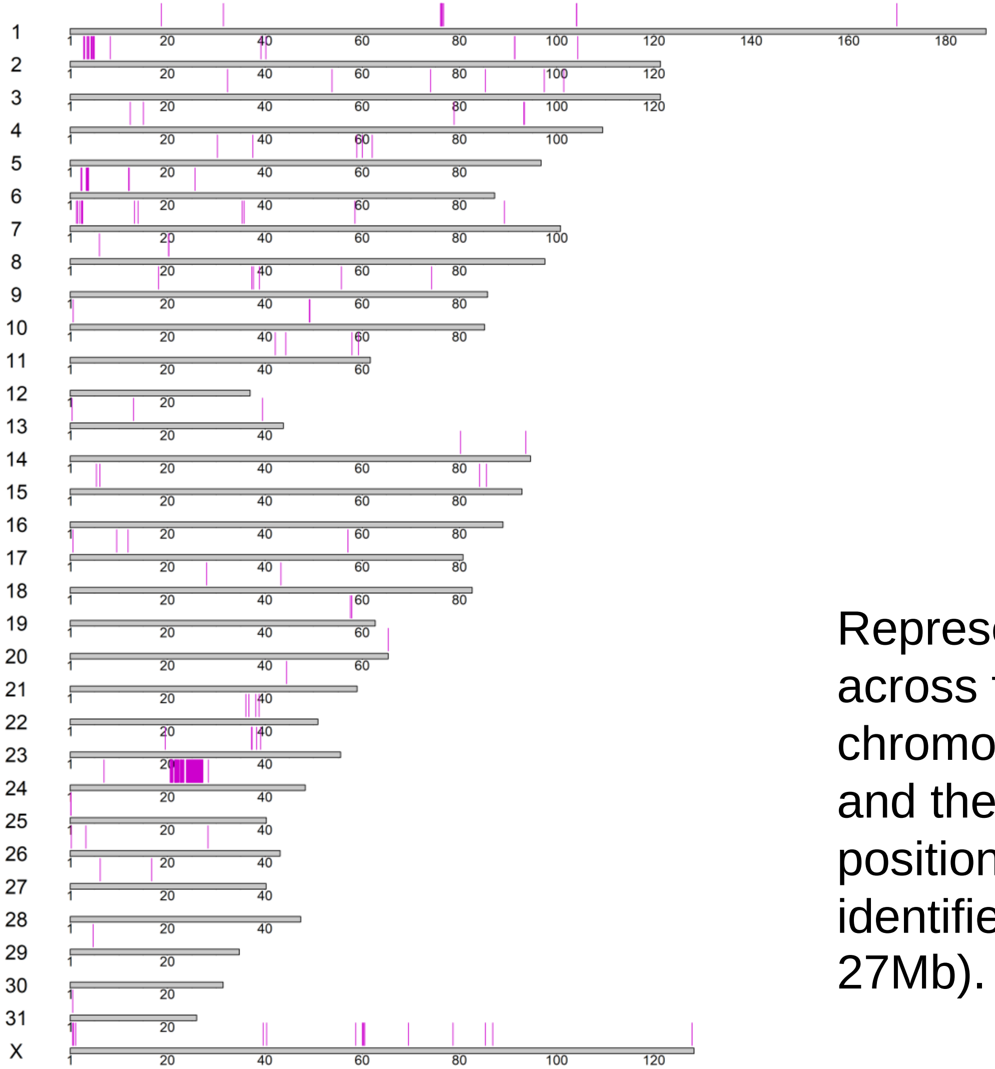


Representation of 3,769 Freebayes variants across the horse genome. Each chromosome is represented by a grey bar and the magenta lines indicate variant positions. A large ~6Mb haploblock was identified on chromosome 24 (21Mb – 27Mb).

**Supplemental table 3.** Associated Variants with P<0.0001

| Position | P-values | Nearest Gene | Expression and Interaction in Platelets |
|---|---|---|---|
| chr24:22162998 | 3.74E-05 | *ANGEL1* | No |
| chr24:22642450 | 3.74E-05 | *SAMD15* | No |
| chr24:22642470 | 3.74E-05 | *SAMD15* | No |
| chr24:22888518 | 3.74E-05 | *ALKBH1* | No |
| chr24:26081856 | 7.63E-06 | *SEL1L* | Yes |
| chr24:21867050 | 3.74E-05 | *ESRRB* | No |
| chr24:21999935 | 3.74E-05 | *ESRRB* | No |
| chr24:22685398 | 3.74E-05 | *VIPAR* | Yes |
| chr24:24095086 | 9.73E-06 | *NRXN3* | No |
| chr24:24460139 | 9.73E-06 | *NRXN3* | No |
| chr24:24525130 | 9.73E-06 | *NRXN3* | No |
| chr24:24709312 | 2.29E-05 | *NRXN3* | No |
| chr24:25392066 | 3.74E-05 | *CEP128* | No |
| chr24:25429259 | 3.74E-05 | *CEP128* | No |
| chr24:26447375 | 7.63E-06 | *SEL1L (AL355838.1)* | Yes |

### Supplemental Figure 3

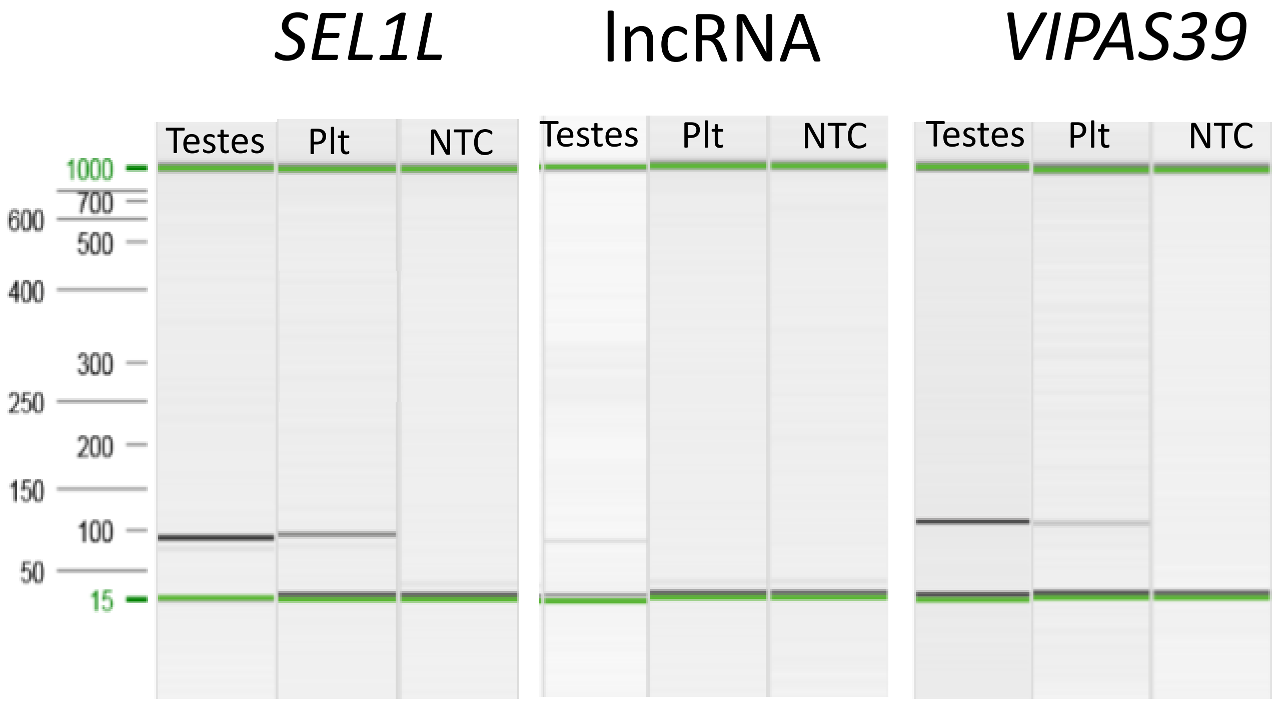


Reverse transcription PCR showing transcript presence in testes (positive control) and platelets (plt). NTC represents no template control. *SEL1L* and *VIPAS39* transcripts are shown to be present in platelets while the lncRNA transcript is not.

### Supplemental Figure 4

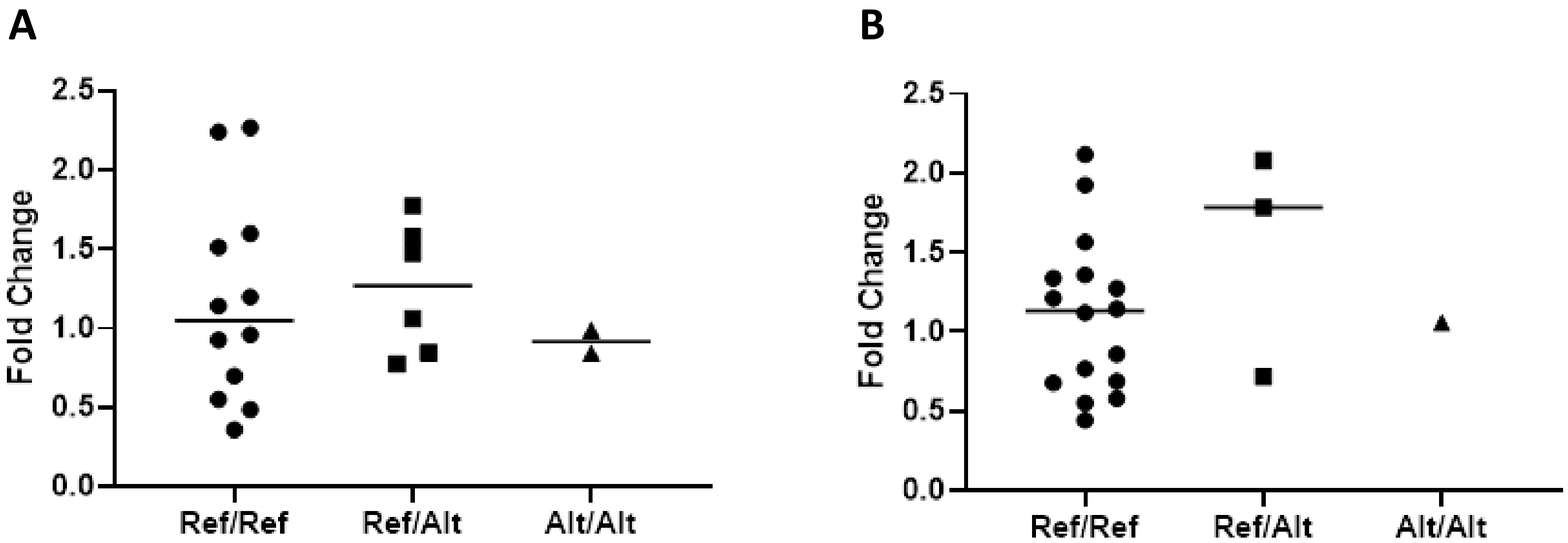


Quantitative reverse transcription PCR from platelet mRNA for (A) *SEL1L* (n=12 Ref/Ref, 6 Ref/Alt, 2 Alt/Alt) (B) *VIPAS39* (n= 16 Ref/Ref, 3 Ref/Alt, 1 Alt/Alt) by genotype. There were no significant differences between genotypes

### Supplemental Figure 5

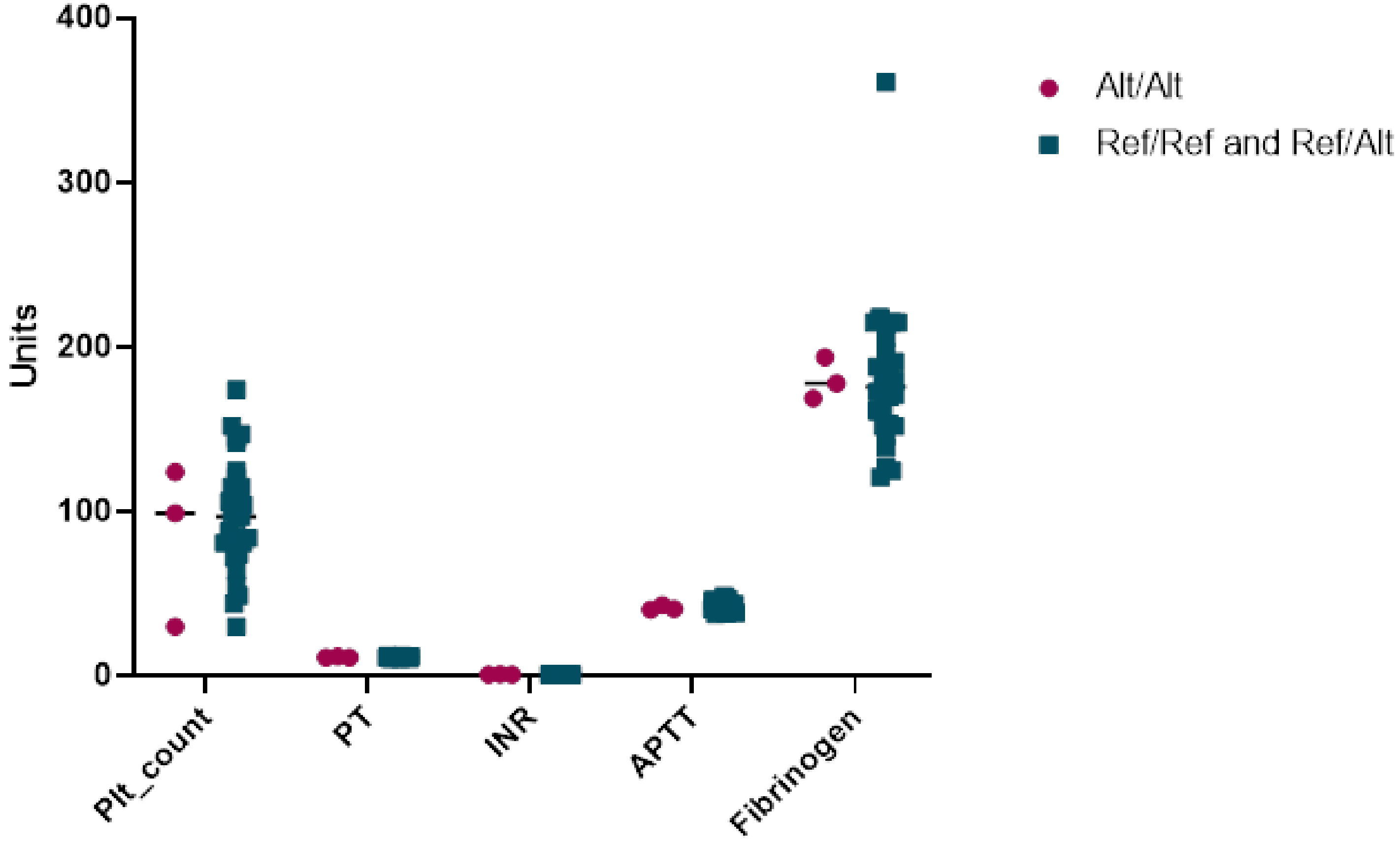


Results of platelet count and coagulation panel indicating that there are no other clotting disorders that may affect the data.

### Supplemental Figure 6

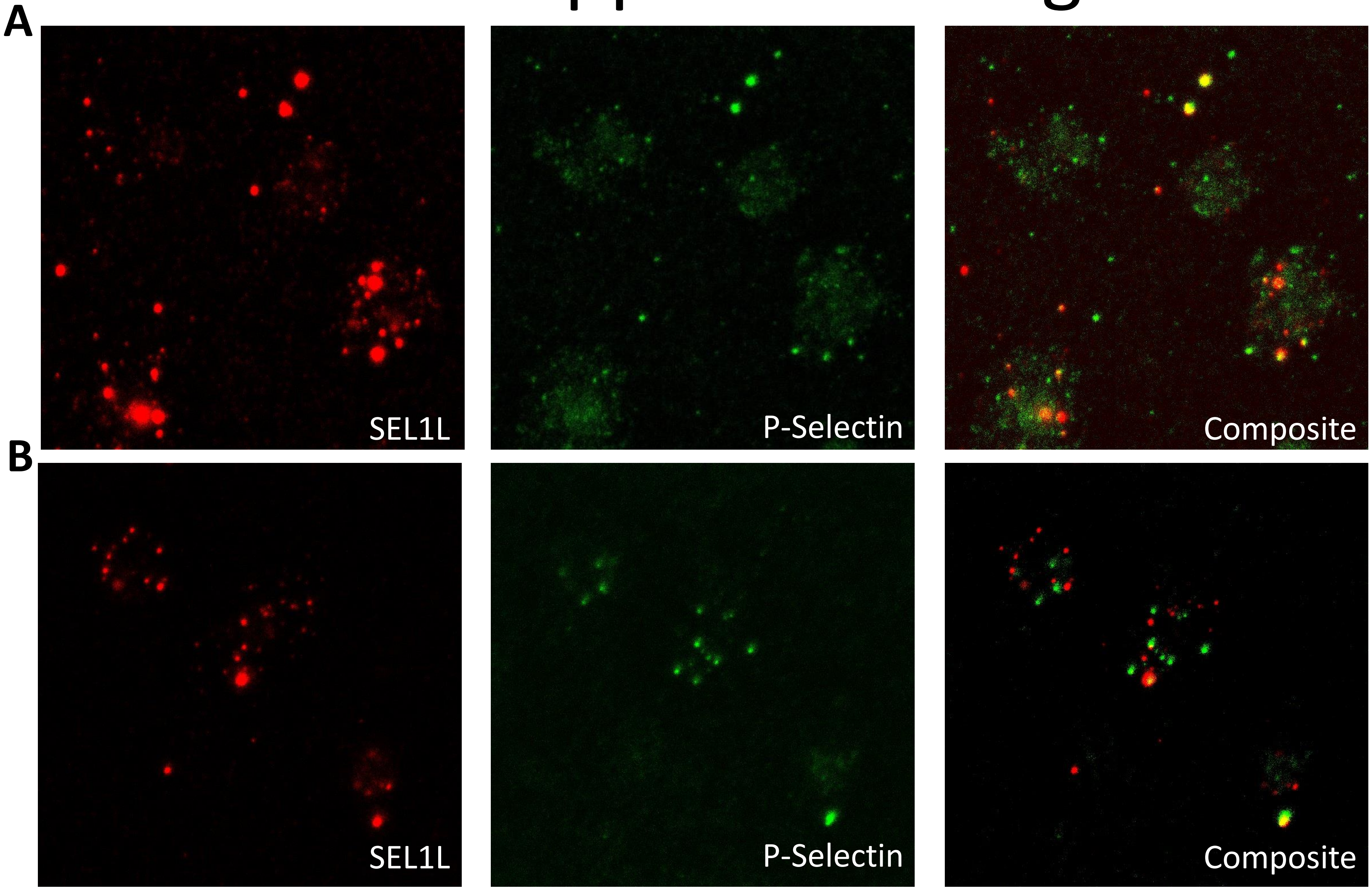


SEL1L (red) localization in relation to P-selectin (green) in (A) resting permeabilized platelets and (B) thrombin activated platelet surface of an Alt/Alt horse. SEL1L did not localize to the surface in resting permeabilized platelets (A) but did localize to the surface in thrombin-activated platelets (B). N=3

### Supplemental Figure 7

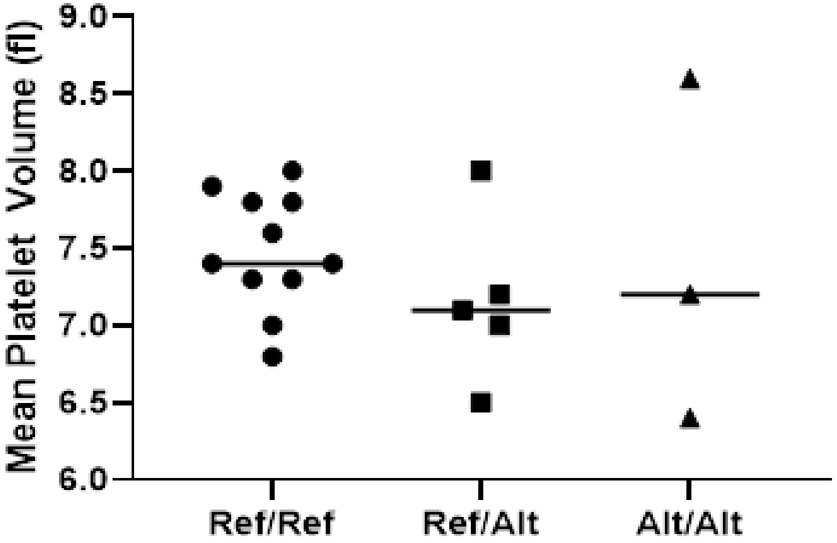


Mean platelet volumes from platelet rich plasma for each individual horse separated by genotype for *SEL1L* c.1810A>G p.Ile604Val. No significant difference was detected among the genotypes. N=11 Ref/Ref, 5 Ref/Alt, 2 Alt/Alt.

### Supplemental Figure 8

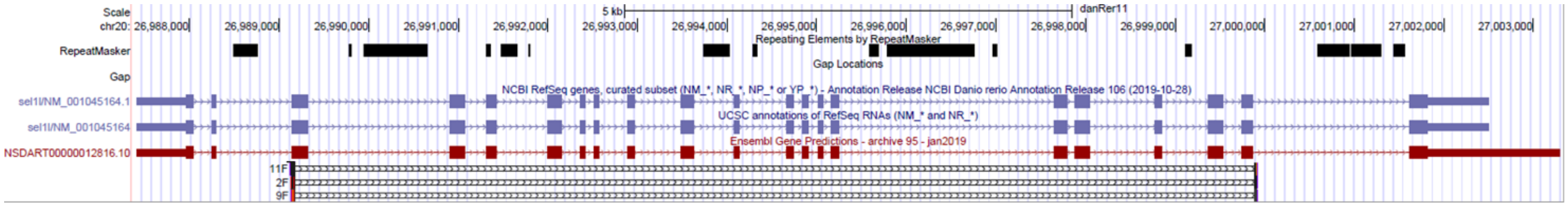


Depiction of CRISPR-Cas9 mediated deletion in *sel1l*. The annotations of the *sel1l* gene are shown in blue and red. The deletion shown spans from exon 3 to exon 20.

### Supplemental Figure 9

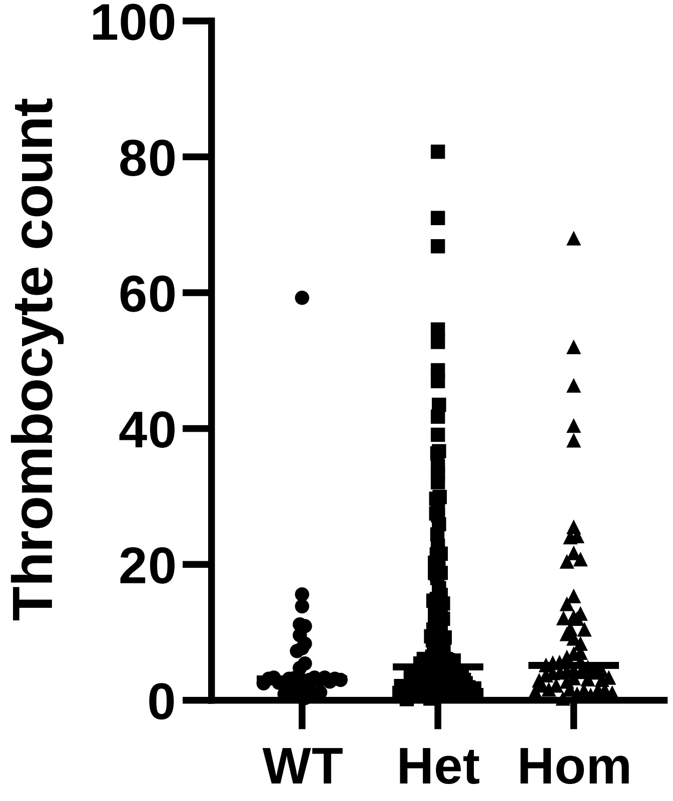


Relative thrombocyte count of wild-type (WT), heterozygous (Het), and homozygous (Hom) offspring from a heterozygous incross of a zebrafish *sel1l* knockout shows no difference in average counts between genotypes. N= 36 WT, 154 Het, 58 Hom. Not significant by ANOVA testing.
